# *foxc1a* and *foxc1b* differentially regulate angiogenesis from arteries and veins by modulating Vascular Endothelial Growth Factor signalling

**DOI:** 10.1101/417931

**Authors:** Zhen Jiang, Teri Evans, Aaron M. Savage, Matthew Loose, Timothy J.A. Chico, Fredericus J.M. van Eeden, Robert N. Wilkinson

**Author notes:** Corresponding author: Robert Wilkinson, Department of Infection, Immunity and Cardiovascular Disease, Medical School, University of Sheffield, S10 2RX, UK.

## Abstract

The forkhead transcription factors *Foxc1* and *Foxc2* are essential to establish intact vascular networks in mammals. How these genes interact with endothelial signalling pathways to exert their functions remains incompletely understood. We have generated novel zebrafish mutants in *foxc1a* and *foxc1b*, the zebrafish orthologues of mammalian *Foxc1*, to determine their function during angiogenesis. *foxc1a* mutants display abnormal formation of cranial veins including the primordial hindbrain channels (PHBC), reduced Vascular Endothelial Growth Factor (VEGF) receptor expression in these and loss of central arteries. *foxc1b* mutants are normal, whereas *foxc1a*; *foxc1b* double mutants exhibit ectopic angiogenesis from trunk segmental arteries. Dll4/Notch signalling is reduced in *foxc1a; foxc1b* double mutant arteries and ectopic angiogenesis can be suppressed by induction of Notch or inhibition of Vegfc signalling. We conclude that *foxc1a* and *foxc1b* play compensatory and context-dependent roles to co-ordinate angiogenesis by promoting venous sprouting via induction of VEGF receptor expression whilst antagonising arterial sprouting by inducing Dll4/Notch signalling. *foxc1a*/*b* mediated induction of both pro- and anti-angiogenic axes of VEGF-Dll4/Notch negative feedback imparts competition to balance arterial and venous angiogenesis within developing vascular beds.

**Summary Statement:** *foxc1a* and *foxc1b* promote angiogenesis from veins and suppress angiogenesis from arteries by promoting competing pro-angiogenic Vascular Endothelial Growth Factor signalling, and anti-angiogenic Dll4/Notch signalling in zebrafish embryos.

## Introduction

The cardiovascular system is the first functional organ system to form during vertebrate embryogenesis and is essential to facilitate growth and development. Within the cardiovascular system, blood vessels represent the primary conduit for nutrient transport and metabolic exchange and these are formed from endothelial progenitor cells, or angioblasts, which are first specified within the lateral plate mesoderm. During somitogenesis, angioblasts differentiate into arterial and venous cell fates and undergo medial migration to coalesce into a solid linear mass of cells or vascular cord. These vascular cords undergo subsequent lumenisation and are remodelled into a functional and complex vascular network by a process of selective cell sprouting termed angiogenesis. Angiogenesis requires complex co-ordination of cell signalling and cellular behaviours to generate a branched vascular morphology capable of maximising surface area for metabolic exchange while minimising transport distances. While angiogenesis is tightly regulated by interactions between pro-angiogenic Vascular Endothelial Growth Factor (VEGF) and anti-angiogenic Notch signalling, reviewed in (Blanco and Gerhardt, 2013), how these pathways are transcriptionally regulated is not fully understood.

VEGF is a morphogen which signals via distinct ligands to induce migratory behaviour within endothelial cells (ECs) and drive blood vessel sprouting. In zebrafish, angiogenesis from arteries is promoted primarily by interaction of Vegfa with its cognate receptor Vegfr4/Kdrl or Vegfr2/Kdr (Bahary et al., 2007; Covassin et al., 2006; Habeck et al., 2002; Nasevicius et al., 2000), whereas Vegfc promotes sprouting from veins via Vegfr3/Flt4 (Hogan et al., 2009b; Le Guen et al., 2014; Villefranc et al., 2013). Within an angiogenic sprout, leading ECs are termed tip cells and these are followed by trailing stalk cells. As an angiogenic sprout forms, a migrating EC extends filopodia to sense VEGF signals and upregulate *Dll4* transcription (Gerhardt et al., 2003; Hellstrom et al., 2007), inducing Notch signalling in neighbouring cells and this acts to limit excessive angiogenic sprouting (Siekmann and Lawson, 2007). Notch signalling inhibits expression of VEGF receptors in neighbouring stalk cells (Lobov et al., 2007), thereby limiting the ability of these cells to respond to VEGF and controlling the number of tip cells per sprout. Flt4 is expressed in angiogenic tip cells (Gore et al., 2011; Shin et al., 2016) and hypersprouting of arteries induced by knockdown of *dll4* can be rescued by inhibition of Vegfc-Flt4 signalling (Hogan et al., 2009b; Villefranc et al., 2013). Thus, the interplay between VEGF and Dll4/Notch signalling dynamically controls behaviour of angiogenic sprouts within developing vascular networks, reviewed in (Blanco and Gerhardt, 2013).

The forkhead family of transcription factors possess a highly conserved forkhead DNA binding domain (Kaufmann and Knochel, 1996; Lai et al., 1993) and more than 40 forkhead box (Fox) proteins have been identified in mammals (Ivanov et al., 2013). Members of the C class of Fox proteins, FOXC1 and FOXC2 have been implicated in vascular development (Kume, 2009) and mutations in *FOXC1* have been described in cerebral small vessel disease which can lead to stroke (French et al., 2014). *FOXC1* is also mutated in Axenfeld-Rieger syndrome which causes craniofacial defects including iris hypoplasia (Smith et al., 2000). Mutations in *FOXC2* cause lymphoedema-distichiasis syndrome and result in primary lymphoedema (Mellor et al., 2011). Murine *Foxc1* or *Foxc2* mutants exhibit pre-or perinatal lethality with variable cardiovascular and skeletal defects (Iida et al., 1997; Smith et al., 2000; Topczewska et al., 2001a; Winnier et al., 1997; Winnier et al., 1999). *Foxc1; Foxc2* double mutant mice exhibit similar developmental abnormalities to *Foxc1* single mutants but with increased severity and abnormal arteriovenous specification, indicating redundant functions between these genes (Seo et al., 2006; Topczewska et al., 2001a). Murine *Foxc1*; *Foxc2* double mutants die by E9.5 preventing detailed analysis of angiogenesis, however some *Foxc1+/-; Foxc2-/-* embryos survive until E12.5 and exhibit arteriovenous malformations (Seo et al., 2006).

In teleosts, *Foxc2* has been lost, while *Foxc1* has undergone duplication to generate the paralagous genes *foxc1a* and *foxc1b* (Topczewska et al., 2001b) and this study. Functional studies of *foxc1a* and *foxc1b* during vascular development have relied on gene knockdown and mutant analysis has been lacking. Knockdown of *foxc1a* and *foxc1b* by morpholino has been reported to induce cerebral haemorrhage and circulation defects (De Val et al., 2008; Skarie and Link, 2009) and this was suggested to arise from arteriovenous malformations and reduced vascular basement integrity (Skarie and Link, 2009). However, reported disruption of angiogenesis induced by knockdown of *foxc1a* and *foxc1b* are inconsistent between studies (De Val et al., 2008; Skarie and Link, 2009). Foxc transcription factors can directly activate the Dll4 and Hey2 promoter in cultured ECs, indicating these genes function to promote Notch signalling (Hayashi and Kume, 2008; Seo et al., 2006). In zebrafish, *foxc1a* synergises with Notch signalling during kidney development and Foxc1a can activate Notch targets *in vitro* (O’Brien et al., 2011), however whether *foxc1a/b* act upstream of Notch during zebrafish vascular development is not known. Foxc2 combinatorically activates EC genes in partnership with Etv2 in Xenopus embryos and *foxc1a/b* knockdown in zebrafish reduced angiogenesis within the developing trunk (De Val et al., 2008). In addition, knockdown of *foxc1a/b* in zebrafish has been reported to reduce Etv2 promoter activity, suggesting these genes also function upstream of Etv2 (Veldman and Lin, 2012). Several recent studies using *foxc1a* mutants have indicated a requirement for *foxc1a* during neural circuit development (Banerjee et al., 2015) in addition to anterior somite formation (Hsu et al., 2015; Li et al., 2015) which is consistent with previous knockdown studies (Topczewska et al., 2001b). However, mutant analysis of *foxc1a* during vascular development and investigation of potential genetic interaction between *foxc1a* and *foxc1b* during this process remain unaddressed.

We have generated novel zebrafish *foxc1a* and *foxc1b* mutants and characterised the function of these transcription factors during angiogenesis. *foxc1a* mutants display abnormal cranial angiogenesis including delayed formation of primordial hindbrain channels and almost total loss of central arteries within the developing hindbrain, whereas *foxc1b* mutants are morphologically normal. By contrast, *foxc1a; foxc1b* double mutants exhibit ectopic sprouting of segmental arteries within the developing trunk and reduced segmental vein formation, in addition to abnormal cranial vessel formation observed in *foxc1a* single mutants. We find that *foxc1a* promotes venous expression of VEGF receptors including *vegfr3/flt4* and *vegfr4/kdrl*, thereby promoting venous angiogenesis, whilst *foxc1a* and *foxc1b* genetically interact to limit angiogenesis from arteries by suppressing Vegfc/Flt4 signalling via induction of Dll4/Notch signalling. Our data indicates *foxc1a* and *foxc1b* play compensatory and context-dependent roles to co-ordinate angiogenesis from arteries and veins via differential regulation of pro-and anti-angiogenic signalling.

## Results

### *foxc1a* mutants display multiple vascular abnormalities while *foxc1b* mutants are morphologically normal

Mammals have two Foxc genes, *Foxc1* and *Foxc2* (Fig. S1, S2), whereas teleost lineages have no identifiable *foxc2* ortholog (Fig. S2) but possess two paralogous *Foxc1* orthologues, *foxc1a* and *foxc1b* (Topczewska et al., 2001b). To investigate the specific functions of *foxc1a* and *foxc1b* during vascular development, we generated zebrafish mutants using genome editing (see methods). Wild type *foxc1a* and *foxc1b* comprise single exon genes, which encode proteins containing 476 and 433 amino acids respectively (Fig. 1A, B) (Topczewska et al., 2001b). The *foxc1a*^*sh356*^ mutant allele contains a 4bp insertion, generating a protein which is predicted to retain the first 56 amino acids of wild type Foxc1a before shifting frame and truncating following 14 incorrect amino acids (Fig. 1A; Fig. S3 A, B). The *foxc1b*^*sh408*^ mutant allele contains a 13bp deletion, which is predicted to shift frame after the first 58 amino acids and prematurely truncate the protein following 10 incorrect amino acids (Fig. 1B; Fig. S3 C, D). Both *foxc1a*^*sh356*^ and *foxc1b*^*sh408*^ alleles are predicted to truncate prior to the conserved Forkhead DNA binding domain (Fig. 1A, B Green box) and are therefore likely to represent severe loss of function or null mutations. These alleles are hereafter referred to as *foxc1a* and *foxc1b*.

**Figure 1.**
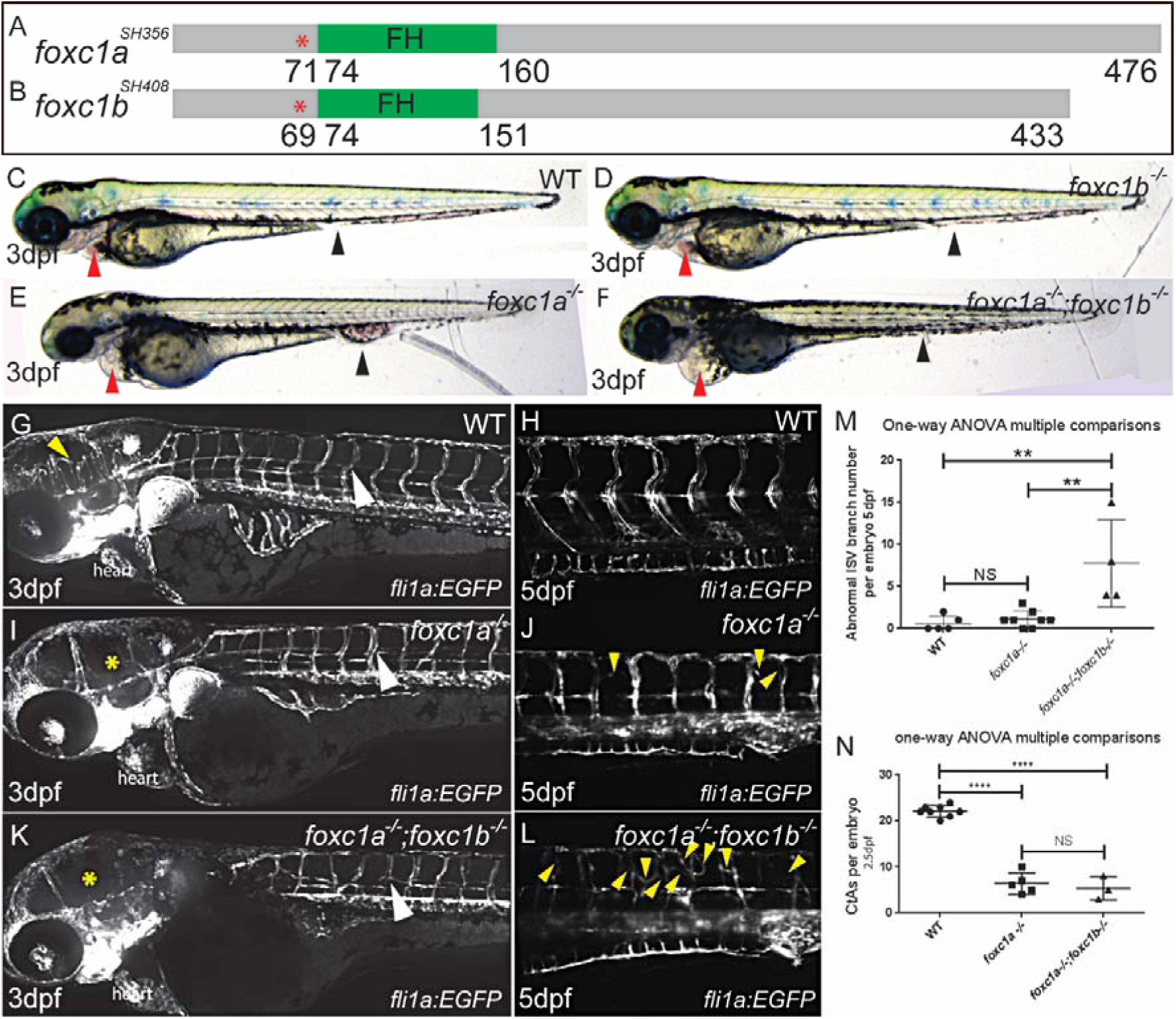
Generation and characterisation of *foxc1a* and *foxc1b* mutant zebrafish. (**A**) Schematic representation of *foxc1a*^*SH356*^ allele and *foxc1b*^*SH408*^ allele (**B**) with premature stop codon after amino acid 70 in *foxc1a* (red asterisk) and amino acid 68 in *foxc1b* (red asterisk). (**C-F**) Morphology of 3dpf WT sibling (**C**), *foxc1b*^*-/-*^ (D), *foxc1a*^*-/-*^ (E), and *foxc1a*^*-/-*^; *foxc1b*^*-/-*^ double mutants (**F**). Red arrowheads highlight presence of pericardial oedema; black arrowheads highlight region of caudal blood pooling. (**G**, **I**, **K**) Confocal microscopy of 3dpf *Tg(fli1a:EGFP)* WT sibling, *foxc1a*^*-/-*^ and *foxc1a*^*-/-*^;*foxc1b*^*-/-*^double mutants. Yellow arrowhead indicates CtAs in WT (**G**), yellow asterisks indicate absent CtAs in *foxc1a*^*-/-*^ (**I**) and *foxc1a*^*-/-*^; *foxc1b*^*-/-*^ double mutants (**K**); white arrowheads denote ISVs. (**H**, **J**, **L**) Light-sheet microscopy of 5dpf *Tg(fli1a:EGFP)* WT sibling, *foxc1a*^*-/-*^ and *foxc1a*^*-/-*^; *foxc1b*^*-/-*^ double mutants. Yellow arrowheads indicate abnormal ISV sprouts. (**M**) Numbers of ectopic ISV sprouts per embryo at day 5. One-way ANOVA multiple comparisons. **<0.01, NS: not significant. (N) Number of CtAs per embryo at 2.5dpf. One-way ANOVA multiple comparisons. ****<0.0001, NS: not significant. CtA: Central Artery; FH, Forkhead DNA binding domain; ISV, Intersegmental Vessel; LDA, Lateral Dorsal Aorta; PHBC, Primordial Hindbrain Channel; WT, wild-type.

*foxc1a* single mutants displayed absent or substantially reduced blood circulation throughout the developing trunk, variable pooling of blood within the caudal vein plexus (Fig. 1E black arrowheads) and severe pericardial oedema in comparison to wild type siblings (Fig. 1C, E red arrowheads). Some *foxc1a* mutants retained the cranial circulatory loop comprised of the heart, ventral aorta, lateral dorsal aortae, primitive internal carotid artery, basilar artery, posterior hindbrain channel and common cardinal vein (not shown). *foxc1a; foxc1b* double mutants were morphologically similar to *foxc1a* single mutants (Fig. 1F) and displayed similar circulation abnormalities. Most *foxc1a* and *foxc1a; foxc1b* mutants died before 6dpf whereas *foxc1b* mutants were morphologically normal (Fig. 1D), displayed normal circulation and were homozygous viable and fertile (not shown).

No observable vascular defects were present in *foxc1b* mutants (not shown), however, *foxc1a* mutants displayed straightening of intersegmental vessels (ISVs) by 3dpf (Fig. 1I white arrowhead, J yellow arrowheads) in comparison to the normal chevron shape illustrated by ISVs in WT sibs (Fig. 1G white arrowhead, H) as these follow vertical myotomal boundaries. Previous studies have demonstrated functions for *foxc1a* during somite formation and patterning (Hsu et al., 2015; Li et al., 2015; Topczewska et al., 2001b) and abnormal somite formation is known to influence the migratory path of ECs (Shaw et al., 2006). We therefore examined somite morphology in our *foxc1* mutants and identified that while somite morphology was normal in *foxc1b* mutants consistent with normal ISV patterning in these (Fig. S4B arrowheads), the formation of somites 1-4 was abnormal in *foxc1a* mutants (Fig. S4A, C, E, F red arrowheads, G) consistent with previous reports (Hsu et al., 2015; Li et al., 2015). In addition, *foxc1a* mutants displayed a significant increase in somitic angle of posterior somites (Fig S4E, F, H), which likely accounts for straightened ISVs in these mutants. No difference in somite morphology were observed between *foxc1a* mutants and *foxc1a; foxc1b* double mutants (Fig. S4C, D). While ISV formation in *foxc1a* mutants was unremarkable (Fig. 1H, J arrowheads, M), *foxc1a; foxc1b* double mutants surprisingly displayed a significant increase in frequency of misbranched ISVs within the developing trunk at 5dpf (Fig. 1J, L arrowheads, M). By contrast, both *foxc1a* and *foxc1a;foxc1b* double mutants exhibited significantly fewer central arteries (CtAs) within the developing brain at 3dpf in comparison to WT (Fig. 1G, I, K, yellow arrowhead and asterisks, N). Strikingly, these data suggest *foxc1a* and *foxc1b* differentially regulate angiogenesis within the developing zebrafish trunk and brain.

### *foxc1a* is required for central artery formation

Since blood circulation was absent throughout the trunk in the majority of *foxc1a* and *foxc1a; foxc1b* double mutants but cranial circulation was retained in some embryos, we first examined the hindbrain region where these two circulatory loops interconnect, using confocal time lapse microscopy (Fig. S5). *foxc1a* mutant embryos displayed delayed angioblast migration during the formation of primordial hindbrain channels (PHBC) (Fig. S5B, Red asterisks). Formation of PHBCs and the basilar artery (BA) were delayed by approximately 3 hours in *foxc1a* mutants (Fig. 2 A-B’, Supplementary Movies 1 and 2) and increasingly delayed in *foxc1a; foxc1b* double mutants by approximately 6 hours (Fig. S5C) Formation of common cardinal veins (CCV) (Fig. S6A-D) and lateral dorsal aortae (LDA) (Fig. S6E-G) were also abnormal in *foxc1a* mutants. In addition, angiogenic sprouting of CtA was greatly reduced in *foxc1a* mutants at 60hpf (Fig. 2A’, B’ C-F arrowheads, K, L, Supplementary Movie 2). *foxc1a* mutants also displayed significantly increased frequency of arterial venous connections (AVCs) between the PHBC and the BA or posterior communicating segments (PCS) (Fig. 2M). EC number within basal cranial vessels (PHBC, BA, PCS) were not significantly altered in *foxc1a* mutants (Fig. 2N), indicating that reduced CtA formation was not due to reduced EC number in basal cranial vessels. Overexpression of full length *foxc1a* mRNA partially rescued CtA (Fig. 2C-L, arrowheads) and AVC frequency (Fig. 2L, M) in *foxc1a* mutants, confirming that loss of these vessels was due to mutation of *foxc1a*.

**Fig 2.**
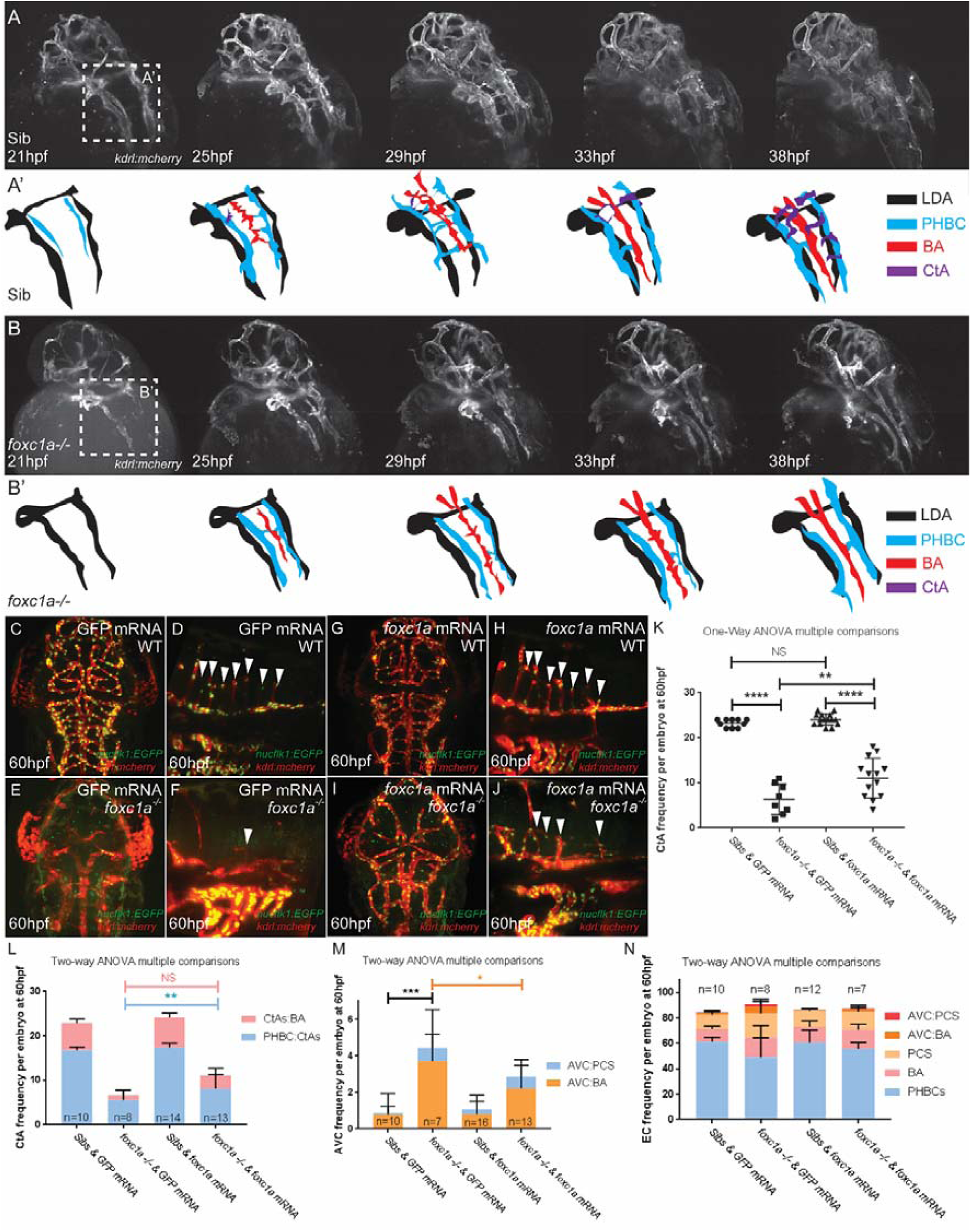
*foxc1a* is required for central artery formation. **A**, **B**) Lightsheet time lapse images and corresponding illustrations of cranial vessel formation in *Tg*(*kdrl:mcherry*) sibling (**A, A’**) and *foxc1a* mutant embryo (**B**, **B’**) between 21hpf and 38hpf. *foxc1a* mutants (**B, B’**) exhibit delayed PHBC formation and reduced CtA formation. Cranial vessels are colour coded (**A’**, **B’)**. **C-K**) Dorsal and lateral views of representative embryo are shown. Control embryos injected with 700pg GFP show normal CtA formation in *Tg(kdrl:mcherry; flk1:nlsEGFP) sibs* (**C**, **D**, arrowheads, **K**) and reduced CtA formation in *foxc1a* mutants (**E**, **F**, arrowheads, **G,H**) *foxc1a* mRNA overexpression is not sufficient to perturb normal central artery formation (**G**, **H**, arrowheads, **K**). **I**, **J**) *foxc1a* mutants injected with 700pg *foxc1a* mRNA display a significant increase in CtA frequency (**I**, **J**, arrowheads, **K**) in comparison to controls (**E**, **F**, arrowheads, (**K**, each data point represents an embryo. One-way ANOVA multiple comparisons, ****<0.0001, **<0.01). **L**) Quantification of CtAs distribution per embryo at 60hpf. Two-Way ANOVA multiple comparisons, **<0.001, NS: not significant. CtAs:BA, CtAs to BA connection; PHBC:CtAs, PHBC to CtAs connection. **M**) Quantification of arterial-venous connection frequency. Two-way ANOVA multiple comparisons. *<0.05, ***<0.001. AVC:PCS, AVC between PHBC and PCS; AVC:BA, AVC between PHBC and BA. **N**) Quantification of endothelial cell number in cranial basal vessels at 60hpf. Two-way ANOVA multiple comparisons. AVC:PCS, EC numbers in AVC between PHBC and PCS; AVC:BA, EC numbers in AVC between PHBC and BA. AVC, Arterial Venous Connection; BA, Basal Artery; CtA, Central Artery; EC, Endothelial cell; PHBC, Primordial Hindbrain Channel; PCS, Posterior communicating Segments.

### *foxc1a* mutants display reduced expression of genes required for normal formation of PHBCs and CtAs

Delayed PHBC formation in *foxc1a* mutants (Fig. 2, S5) is very similar to that induced by *vegfr3/flt4* and *vegfc* loss of function (Covassin et al., 2006; Hogan et al., 2009b; Shin et al., 2016; Villefranc et al., 2013). By contrast, while Vegfc signalling through Flt4 is dispensable for CtA formation (Hogan et al., 2009b; Le Guen et al., 2014), the VEGF receptor *kdrl* is required for CtA sprouting from the PHBC but is dispensable for PHBC formation (Bussmann et al., 2011; Habeck et al., 2002). Consistent with this, expression of *flt4* was reduced within cranial vessels including the PHBC (Fig. 3A, B black arrowheads) and CCV (Fig. 3A, red arrowheads, B, red asterisks). Furthermore, persistent reductions in *flt4* expression were observed in the PHBC in *foxc1a* mutants after it had formed (Fig. 3C, D black arrowheads). *kdrl* expression was also substantially reduced in cranial vessels including the PHBC (Fig. 3E, F black arrowheads) and CCV (Fig. 3E red arrowheads, F red asterisks) in *foxc1a* mutants, which would explain lack of CtAs. In addition, expression of *sox7*, a transcription factor required for formation of central arteries (Hermkens et al., 2015), was substantially reduced in cranial vessels including the PHBC (Fig. 3G black arrowhead, H asterisk) and the CCV (Fig. 3G’ red arrowheads, H’ red asterisks). Taken together, these data indicate *foxc1a* contributes to expression of genes which control formation and sprouting of the PHBC and suggest reduced expression of these in *foxc1a* mutants induce defective PHBC and CtA formation.

**Figure 3.**
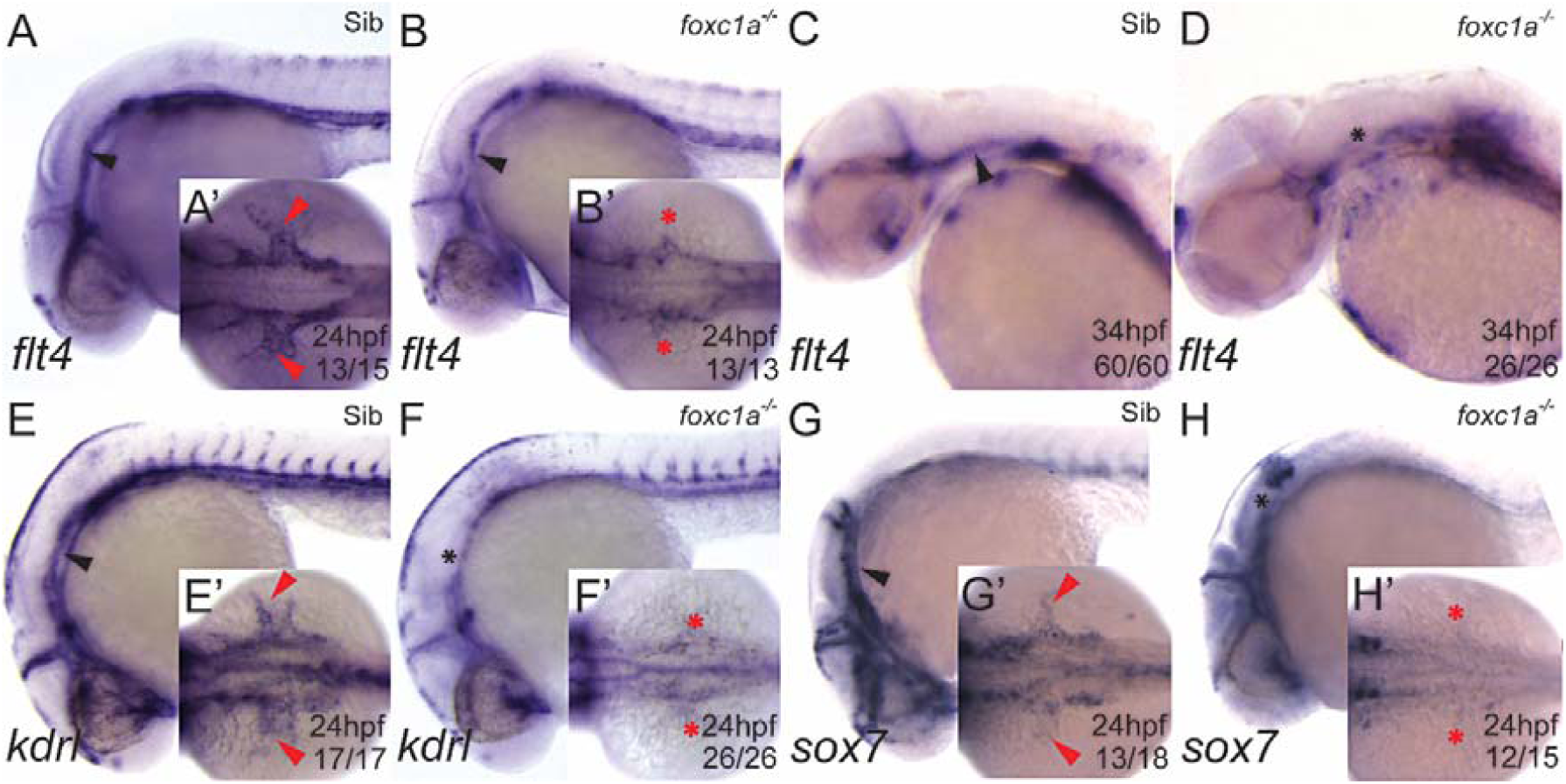
*foxc1a* mutants display reduced expression of genes required for normal formation of primordial hindbrain channels and central arteries. **A-D**) Lateral view of *flt4* expression in WT siblings (**A**, **C**) and *foxc1a* single mutants (**B**, **D**) at 24hpf and 34hpf. **A**’, **B**’) Dorsal view of embryo shown in corresponding panel. Black arrowheads indicate normal or reduced expression of *flt4* in PHBC, respectively; red arrowheads or asterisks denote normal or missing CCV expression, respectively. **E**-**H**) Lateral view of *kdrl* (**E**, **F**) and *sox7* (**G**, **H**) expression in WT sibling (**E**, **G**) and *foxc1a* single mutant (**F**, **H**) at 24hpf. **E’**-**H’**) Dorsal view of embryo shown in corresponding panel. Red arrowheads or asterisks denote normal or missing CCV expression, respectively.

Previous studies have reported expression of *foxc1a* within cranial mesenchyme and hyaloid vasculature (Skarie and Link, 2009). Since *foxc1a* mutants lacked central arteries (Fig. 2) and expression of genes which co-ordinate their formation (Fig. 3), we examined whether *foxc1a* was expressed within PHBCs using fluorescent *in situ* hybridisation and immunocytochemistry (Fig. S7). *foxc1a* was expressed widely throughout cranial mesenchyme and the developing eye (Fig. S7A, A”) as previously reported (Skarie and Link, 2009). *foxc1a* also co-localised with the endothelial reporter *fli1:EGFP* in the mid cerebral vein (MCeV) (Fig S7. A-B”, white arrowheads), CCV (Fig S7. A-B”, white arrowheads), LDA (Fig S7. A-B”, red arrowheads) and PHBC (Fig S7. A-C”, blue arrowheads) at 28hpf and PHBC, CCV and LDA formation are abnormal in *foxc1a* mutants (Fig. S5, S6). Interestingly, while *foxc1a* was clearly expressed within ECs of cranial blood vessels and co-localised with an endothelial marker in the PHBC, CCV and LDA *foxc1b* expression was excluded from these (Fig. S7 D-F”, arrowheads) and displayed perivascular expression (Fig. S7F-F”, blue arrowheads).

### *foxc1a; foxc1b* double mutants display ectopic arterial angiogenesis and reduced venous angiogenesis

Whereas ISVs were straightened in *foxc1a* mutants, their patterning within the developing trunk was relatively normal (Fig. 1I, J, arrowheads). By contrast, *foxc1a; foxc1b* double mutants displayed ectopic ISV branching (Fig. 1L, yellow arrowheads, M) suggesting *foxc1a* and *foxc1b* interact genetically and negatively regulate angiogenesis in this region. Ectopic ISV sprouting in *foxc1a*; *foxc1b* double mutants was observed as early as 28hpf (Fig. 4A-C) and since secondary angiogenesis originating from the posterior cardinal vein (PCV) does not begin until 32hpf (Yaniv et al., 2006) this suggests ectopic sprouts were arterial. Consistent with this, the majority of ectopic vessels within the trunk of *foxc1a*; *foxc1b* double mutants at 4dpf expressed the arterial marker *Tg*(*0.8flt1:RFP*), hereafter referred to as *Tg*(*flt1:RFP*) (Bussmann et al., 2010), (Fig. 4D-F’’’ red arrowheads), indicating these were ectopic segmental arteries (SeA). At 28hpf, *foxc1a* was expressed within the dorsal aorta (DA), SeAs, PCV and surrounding tissue (Fig. S8A-C), however *foxc1b* expression was excluded from trunk ECs at this stage (Fig. S8D-F).

**Figure 4.**
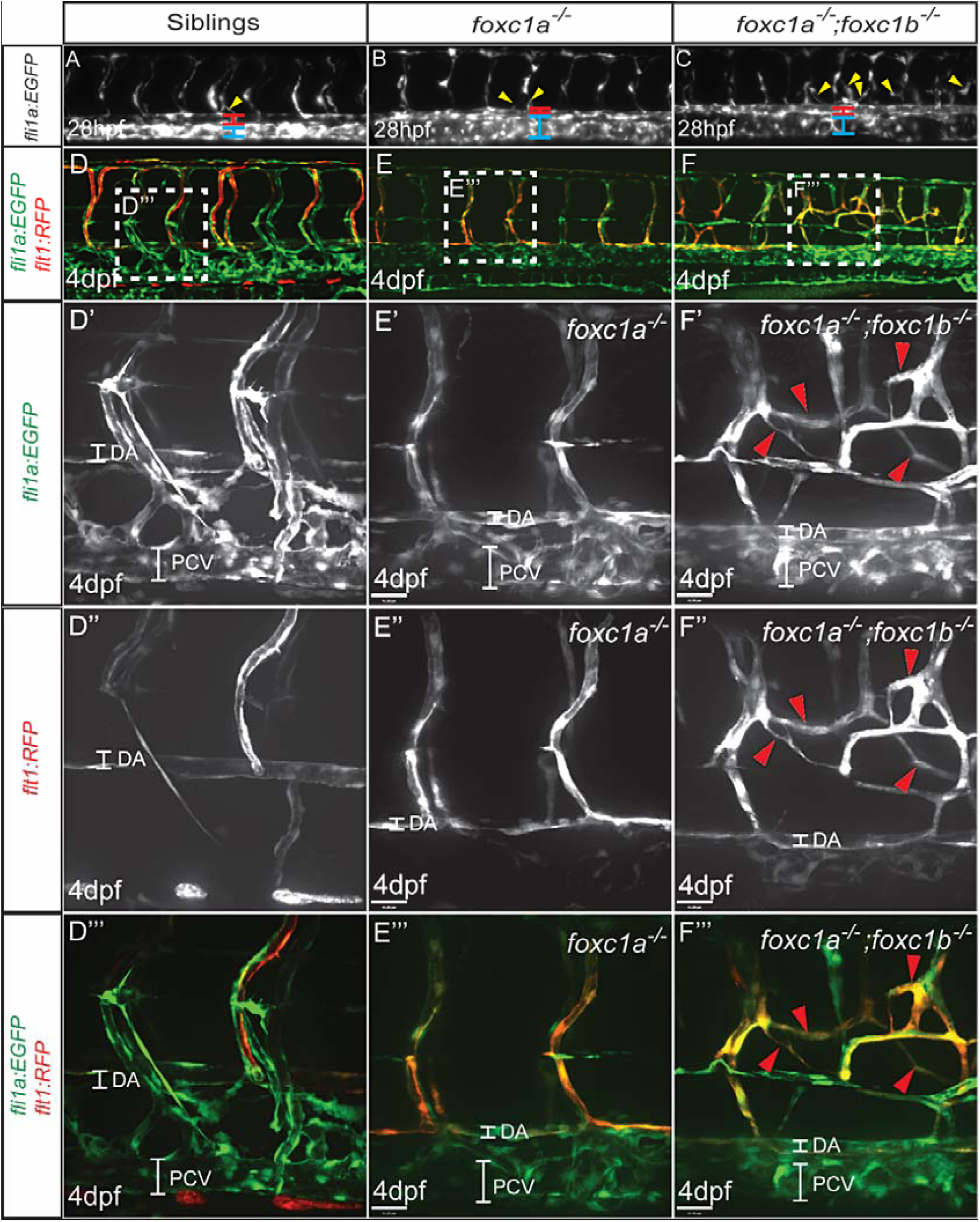
*foxc1a; foxc1b* double mutants display ectopic arterial angiogenesis. **A-C**) Lateral view of trunk vasculature in representative WT sib (**A**), *foxc1a* single mutant (**B**) and *foxc1a; foxc1b* double mutant (**C**) in *Tg*(*fli1a:EGFP*) background at 28hpf. ISV misbranching events are highlighted with yellow arrowheads. **D-F**) Lateral view of trunk vasculature in representative WT sib (**D**), *foxc1a* single mutant (**E**) and *foxc1a; foxc1b* double mutant (**F**) in *Tg*(*fli1a:EGFP;flt1:RFP*) at 4dpf. Highlighted area in D-F corresponds to (**D’, E’, F’**) of the *Tg*(*fli1a:EGFP*) transgene (green channel) shown at increased magnification highlighting ectopic blood vessel formation in *foxc1a; foxc1b* double mutants (F’, red arrowheads). **D’’, E’’, F’’**) Increased magnification of highlighted area displaying the *Tg*(*flt1:RFP*) transgene (red channel). Ectopic segmental arteries in *foxc1a; foxc1b* double mutants are highlighted (F” red arrowheads) **D’’’, E’’’, F’’’**) Merged red and green channel of D’-F” highlighting ectopic segmental artery formation in *foxc1a; foxc1b* double mutants (F’’, red arrowheads). ISV, Intersegmental Vessel; SeA, Segmental Artery; DA, Dorsal Aorta; PCV, Posterior Cardinal Vein.

Since arterial angiogenesis was increased in *foxc1a; foxc1b* double mutants while cranial vein formation was inhibited in both *foxc1a* single and *foxc1a; foxc1b* double mutants, we examined segmental vein (SeV) sprouting in these embryos (Fig. S9). Quantification of the relative distribution of SeAs and SeVs in *foxc1a* and *foxc1a; foxc1b* double mutants using a *Tg*(*fli1a:EGFP;flt1:RFP*) background, revealed a 50% reduction in SeV frequency in both mutants (Fig S9A). Since *flt4* expression was reduced in cranial veins of *foxc1a* mutants (Fig. 3A-D) and is known to promote SeV sprouting from the PCV (Hogan et al., 2009b), we examined its expression within the developing trunk vasculature. From 48hpf, *flt4* expression was also reduced in SeV sprouts of *foxc1a* and *foxc1a; foxc1b* double mutants at 48hpf (Fig. S9B-D, green arrowheads). Collectively, these data indicate venous angiogenesis is impaired in the developing trunk of *foxc1a* and *foxc1a*; *foxc1b* double mutants and suggest *foxc1a* and *foxc1b* positively regulate expression of *flt4* throughout the developing vasculature.

VEGF signalling is essential for SeA formation in zebrafish (Bahary et al., 2007; Covassin et al., 2006; Habeck et al., 2002; Lawson et al., 2003; Nasevicius et al., 2000; Rossi et al., 2016; Shin et al., 2016; Weinstein and Lawson, 2002) and VEGF receptor expression was reduced in trunk (Fig. S9) and cranial (Fig. 3) vessels of *foxc1a* mutants. We therefore examined expression of additional components of VEGF signalling in *foxc1a* and *foxc1a; foxc1b* double mutants (Fig. 5). Vegfa is the major VEGF isoform which promotes SeA formation in zebrafish and *vegfaa* expression was normal in *foxc1a* and *foxc1a*; *foxc1b* double mutants (Fig. 5A-C). However, *kdrl* expression was moderately reduced in the DA (Fig. 5D-F, red arrowheads) and SeAs (Fig. 5D-F, black arrowheads) of *foxc1a* and *foxc1a; foxc1b* double mutants, in keeping with our observations of reduced *kdrl* expression within cranial vessels of *foxc1a* mutants (Fig. 3E, F). By contrast, *flt4*, which is preferentially expressed in venous ECs, was normal within the PCV (Fig. 5 G-I, blue arrowheads), but its expression was not downregulated within SeAs in *foxc1a*; *foxc1b* double mutants by 24hpf in comparison to *foxc1a* mutants (Fig. 5G-I, black arrowheads). Expression of the soluble VEGF decoy receptor, *sflt1* was reduced in the DA (Fig. 5J-K, red arrowheads) and SeAs (Fig. 5J-K, black arrowheads) of *foxc1a* mutants and more substantially reduced in *foxc1a*; *foxc1b* double mutants (Fig. 5L arrowheads). Given that Kdrl promotes SeA formation (Covassin et al., 2006; Habeck et al., 2002) and its expression was not significantly altered between *foxc1a* and *foxc1a*; *foxc1b* double mutants (Fig. 5E, F), we reasoned that the moderate reduction in *kdrl* expression in both mutants compared to wild type sibs was unlikely to account for the ectopic SeA sprouting observed in *foxc1a; foxc1b* double mutants. By contrast, *sflt1* expression was substantially reduced in *foxc1a*; *foxc1b* double mutants compared to *foxc1a* mutants (Fig. 5N, O arrowheads) and this receptor has been shown to antagonise sprouting angiogenesis (Krueger et al., 2011; Zygmunt et al., 2011). We therefore overexpressed *sflt1* in *foxc1a; foxc1b* double mutants and quantified frequency of ectopic SeA sprouts (Fig. S10). Frequency of ectopic SeAs were comparable in *foxc1a*; *foxc1b* double mutants in the presence or absence of *sflt1* overexpression (Fig. S10A-D, arrowheads) indicating reduced *sflt1* expression did not account for ectopic sprout formation in double mutants (Fig. S10E). These data are consistent with recent studies which have demonstrated that *sflt1* limits sprouting from veins, but not arteries (Matsuoka et al., 2016; Wild et al., 2017).

**Figure 5.**
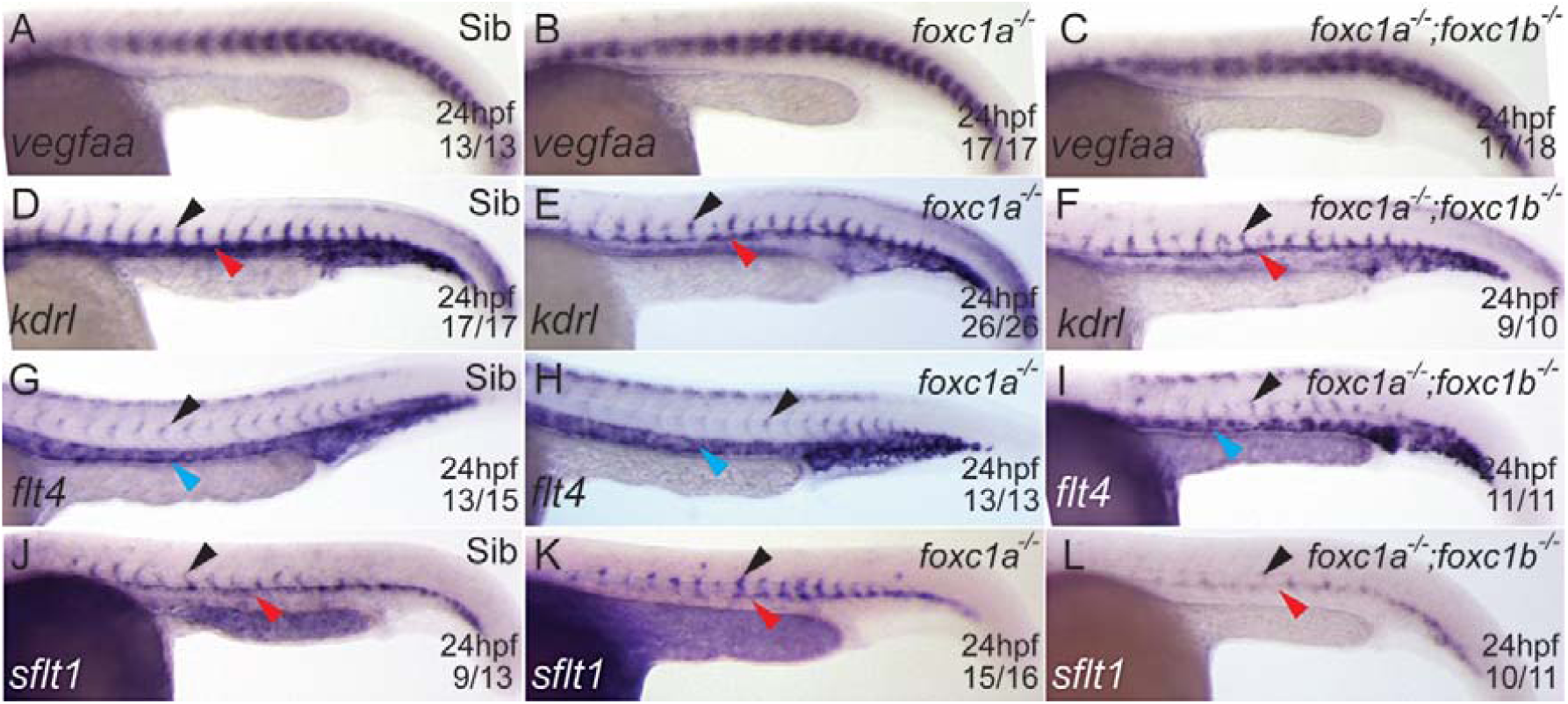
*foxc1a* single mutants and *foxc1a; foxc1b* double mutants display abnormal expression of VEGF receptors within arteries. **A-C**) *In situ* hybridisation for *vegfaa* in siblings, *foxc1a* single mutants and foxc1a; *foxc1b* double mutants at 24 hpf. **D-L**) *In situ* hybridisation for *kdrl* (**D-F**), *flt4* (**G-I**) *sflt1* (**J-L**) in siblings, *foxc1a* mutants and *foxc1a; foxc1b* double mutants at 24hpf. While *vegfaa* expression was normal in *foxc1a* mutants (**A-C**), *kdrl* expression was reduced in both mutants (**D-F**) *flt4* expression was normal in *foxc1a* mutants but was retained in SeAs in *foxc1a; foxc1b* double mutants (**H, I**, black arrowheads). *sflt1* expression was reduced in both mutants (**J-L** arrowheads). Representative embryos are shown. Red arrowheads indicate DA; black arrowheads denote SeA; Blue arrowheads indicate PCV expression.

### *foxc1a and foxc1b* negatively regulate arterial angiogenesis by promoting Notch dependent suppression of *vegfc/flt4* signalling

Foxc1 directly activates the promoters of the Notch ligand Dll4 and Notch target Hey2 *in vitro* (Hayashi and Kume, 2008). Dll4/Notch signalling limits angiogenic sprouting from arteries *in vivo* (Geudens et al., 2010; Leslie et al., 2007; Siekmann and Lawson, 2007) and suppresses Vegfc/Flt4 signalling in SeAs (Hogan et al., 2009b). Thus a loss of Dll4/Notch signalling in *foxc1a; foxc1b* mutants might explain the ectopic sprouting. We therefore examined whether Dll4/Notch signalling was disrupted in these mutants. Activity of a *dll4* enhancer (Sacilotto et al., 2013) was substantially reduced in developing arteries of *foxc1a* mutants and was further reduced in *foxc1a*; *foxc1b* double mutant SeAs (Fig. 6A-C, black arrowheads) and DA (Fig. 6A-C, red arrowheads). Furthermore, expression of *dll4* (Fig. 6D-F) and the Notch target *hey2/gridlock* (Fig. 6G-I) were substantially reduced in *foxc1a; foxc1b* double mutant SeAs (Fig. 6D-I, black arrowheads) and DA (Fig. 6D-I, red arrowheads), indicating Dll4/Notch signalling was reduced in arteries. To determine if ectopic sprouting in *foxc1a*; *foxc1b* double mutants was Notch dependent we crossed *Tg(hs:gal4); Tg(5xUAS-E1b:6xMYC-notch1a)* (Scheer and Campos-Ortega, 1999), into the *foxc1a/+; foxc1b/+* background and subjected progeny to heat shock between 18-20 somite stage to induce expression of the Notch intracellular domain. Heat shock induction of NICD substantially reduced ectopic SeA sprouting in *foxc1a*; *foxc1b* double mutants, thereby demonstrating increased arterial angiogenesis in *foxc1a*; *foxc1b* double mutants was due to reduced Notch signalling.

**Figure 6.**
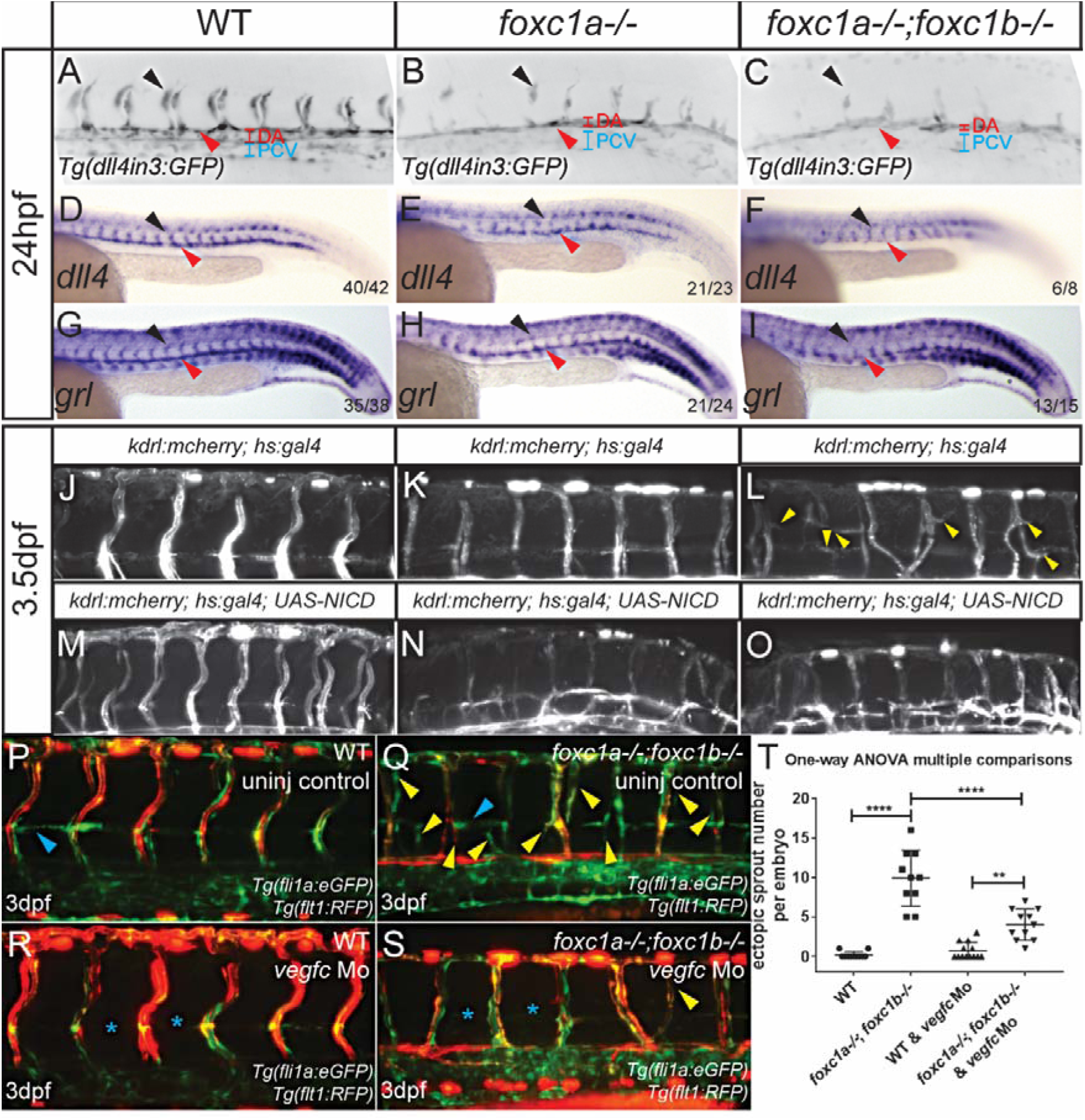
*foxc1a* mutants and *foxc1a; foxc1b* double mutants display reduced Dll4/Notch activity in arteries and ectopic SeA formation can be rescued by Notch induction or *vegfc* knockdown in *foxc1a*; *foxc1b* double mutants. **A-C**) Arterial GFP expression in *Tg*(*dll4in3:GFP*) was reduced in *foxc1a* single mutants (**B**) and *foxc1a; foxc1b* double mutants (**C**) in comparison to sibs (**A**) at 24hpf. Black arrowheads denote SeAs, red arrowheads denote dorsal aorta. **D-I**) *In situ* hybridisation for *dll4* (**D-F**), *grl* (**G-I**) in siblings, *foxc1a* mutants and *foxc1a; foxc1b* double mutants at 24hpf. *dll4* and *grl* expression was reduced in *foxc1a* mutants (**E**, **H**) and substantially reduced in *foxc1a; foxc1b* double mutants (**F**, **I**), Black arrowheads denote SeAs, red arrowheads denote dorsal aorta. **J-O**) Light sheet images of trunk vasculature at 3.5dpf in heat shocked WT sibs (**J**), *foxc1a* single mutants (**K**) and *foxc1a; foxc1b* double mutants (**L**) in *Tg(kdrl:mcherry; hs:gal4)* background and *Tg(kdrl:mcherry; hs:gal4; UAS-NICD)* background in WT sibs (**M**), *foxc1a* single mutants (**N**) and *foxc1a; foxc1b* double mutants (**O**). Yellow arrowheads point to ectopic SeAs (**L**), which are rescued by Notch induction in *foxc1a; foxc1b* double mutants (**O**). **P-S**) Light sheet images of trunk vasculature at 3dpf in *Tg(fli1a:eGFP;flt1RFP)* background. Injection of *vegfc* morpholino suppresses ectopic SeA formation in *foxc1a*; *foxc1b* double mutants (**Q, S**). Blue arrowheads or asterisks denote normal and absent parachordal lymphangioblasts; yellow arrowheads indicate ectopic SeA. (**T**) Quantification of ectopic SeA number in control and *vegfc* Mo injected groups at 3dpf. One-way ANOVA multiple comparisons. **<0.01, ****<0.0001. SeA, segmental artery.

Dll4/Notch signalling suppresses Vegfc/Flt4 signalling in SeAs (Hogan et al., 2009b). Since ectopic SeA sprouting in *foxc1a; foxc1b* double mutants was Notch dependent (Fig. 6J-O) and *flt4* expression was not retained in SeAs in these mutants (Fig. 5I, black arrowhead), we hypothesised that reduced Dll4/Notch signalling in *foxc1a; foxc1b* double mutant arteries may induce ectopic SeA angiogenesis by enhanced Vegfc/Flt4 signalling. We therefore injected *vegfc* morpholino (Hogan et al., 2009a) into *foxc1a; foxc1b* double mutant embryos and quantified frequency of ectopic SeAs (Fig. 6P-T). We observed reduced formation of parachordal lymphangioblasts in *vegfc* morphants (Fig. 6R, S, asterisks) as previously described (Hogan et al., 2009a; Hogan et al., 2009b) indicating Vegfc function was inhibited. The frequency of ectopic SeAs in *foxc1a; foxc1b* double mutants was significantly reduced by *vegfc* knockdown in comparison to control double mutants (Fig. 6Q, S, T, arrowheads). Collectively, our data indicates *foxc1a* and *foxc1b* antagonise angiogenesis from arteries by promoting Dll4/Notch-mediated repression of Vegfc/Flt4 signalling. Conversely, *foxc1a* also promotes venous angiogenesis by positively regulating VEGF receptor expression in veins (Fig. 7). Thus, *foxc1a* and *foxc1b* act in concert to balance angiogenesis from arteries and veins by promoting context dependent expression of both pro-and anti-angiogenic genes during development.

**Figure 7.**
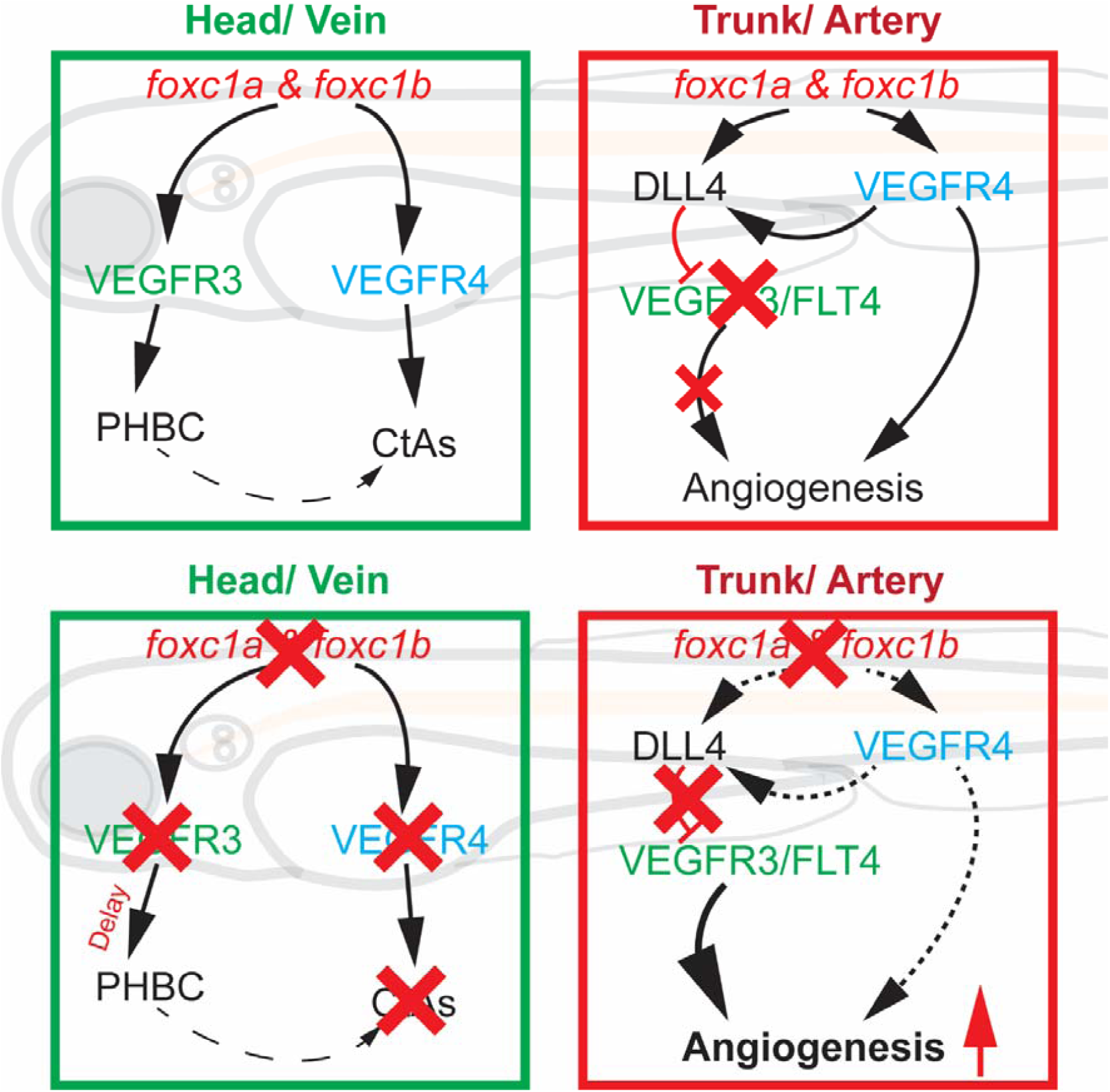
*foxc1a* and *foxc1b* balance angiogenesis by induction of competing pro-and anti-angiogenic signalling

## Discussion

Here we report a detailed genetic analysis of *foxc1a* and *foxc1b* during zebrafish vascular development. Zebrafish *foxc1a* mutants display vascular defects localised to the cranial vasculature, *foxc1a; foxc1b* double mutants display additional defects in trunk vasculature, while *foxc1b* mutant vasculature is normal. This indicates *foxc1a* can fully compensate for loss of *foxc1b* during blood vessel development in zebrafish, whereas *foxc1b* can only partially compensate for loss of *foxc1a* during formation of trunk vasculature. Furthermore, a single copy of *foxc1a* is compatible with normal embryonic development following loss of both *foxc1b* alleles.

Mammals have 2 *Foxc* genes, *Foxc1* and *Foxc2*, however, *Foxc2* has been lost in teleost lineages and *Foxc1* has undergone duplication (Fig. S1, S2). Structural analysis suggests FOXC proteins have the same binding specificity (van Dongen et al., 2000) and are likely to regulate the same downstream targets when co-expressed (Hayashi and Kume, 2008). It therefore seems surprising that *foxc1a* and *foxc1b* mutants have such different phenotypes given that the forkhead DNA binding domain is 97% identical at the protein level in Foxc1a and Foxc1b (Topczewska et al., 2001b) and is probable these transcription factors have highly similar or identical targets. Our data are consistent with this, since we observed greater reductions in expression of the Foxc targets *dll4* and *hey2* (Hayashi and Kume, 2008) in *foxc1a; foxc1b* double mutants than in *foxc1a* mutants. The morphological difference between *foxc1a* and *foxc1b* mutants are therefore not likely to be explained by differential regulation of target genes, but by differential expression of *foxc1a* and *foxc1b* during development, for example, *foxc1a* is expressed in zebrafish ECs whereas *foxc1b* is excluded from these (Fig. S7, S8).

Mutation of *FOXC1* has been proposed to induce human pathologies including cerebral small vessel disease (CSVD) (French et al., 2014). In mice, *Foxc1* is expressed in both pericytes and ECs (Kume et al., 2001; Siegenthaler et al., 2013). Conditional knockout of *Foxc1* in mouse pericytes induces cerebral haemorrhage and as such *Foxc1* function in pericytes has been proposed to maintain integrity of the blood brain barrier in mammals (Siegenthaler et al., 2013). Constitutive *Foxc1* knockout mice exhibit similar defects to pericyte-specific *Foxc1* knockout mice but with much greater phenotypic severity (Kume et al., 1998; Siegenthaler et al., 2013). Interestingly, EC specific inactivation of *Foxc1* in mice does not recapitulate cerebral vascular defects displayed in either pericyte-specific *Foxc1* knockout or constitutive *Foxc1* KO mice (Mishra et al., 2016), indicating *Foxc1* functions non-cell autonomously to promote cerebral blood vessel formation in the mouse. In zebrafish, combined knockdown of *foxc1a* and *foxc1b* by morpholino has been reported to induce cerebral haemorrhage in embryos (French et al., 2014; Skarie and Link, 2009) and this was proposed to occur via reduced vascular basement membrane integrity (Skarie and Link, 2009) or inhibition of *foxc1a*/*foxc1b-*mediated induction of Pdgf signalling in pericytes (French et al., 2014). While we also observe reduced expression of *pdgfrb* in *foxc1a* mutants (not shown) and find both *foxc1a* and *foxc1b* are expressed perivascularly in the zebrafish brain prior to mural cell emergence (Fig. S7), we have not observed cerebral haemorrhage in either *foxc1a* or *foxc1a; foxc1b* double mutant embryos, even in those which retain anterior circulation. These differences could be due to incomplete gene knockdown in morphants, off target effects of morpholinos, or potential compensatory mechanisms which promote vascular integrity in our mutants. However, recent studies in zebrafish have demonstrated that while *pdgfrb* is essential for recruitment of mural cells to cranial vessels including CtAs, mural cells are recruited to cranial vessels such as CtAs only after they have formed and cranial angiogenesis is normal in *pdgfrb* mutants which lack mural cells (Ando et al., 2016). Therefore, since *foxc1a* mutants display reduced cranial angiogenesis, *foxc1a*-mediated induction of *pdgfrb* expression is unlikely to contribute to cranial angiogenesis in zebrafish.

Deletion of *Foxc1* in mouse neural crest-derived cells reproduces cerebrovascular phenotypes of global mouse *Foxc1* mutants (Mishra et al., 2016), indicating its function in this tissue is essential to co-ordinate cranial angiogenesis, however, neural crest cell specification is dispensable for cranial angiogenesis in zebrafish (Ando et al., 2016; Wang et al., 2014). This indicates substantial divergence between *Foxc1*-mediated regulation of vascular development between mouse and zebrafish and suggests that in contrast to mouse, endothelial expression of *foxc1a* is important during cranial angiogenesis in zebrafish. Consistent with this, *foxc1a* is expressed in cranial vessels (Fig. S7), which are abnormal in *foxc1a* mutants while *foxc1b* is excluded from ECs (Fig. S7) and *foxc1b* expression is not induced in ECs in the absence of *foxc1a* (not shown). Our analysis of *foxc1a* and *foxc1b* mutants suggests *foxc1a* is a master regulator of cranial angiogenesis which functions in ECs to promote angiogenesis from cranial veins by inducing VEGF receptor expression and other pro-angiogenic factors including *sox7*. By contrast, *foxc1a* is dispensable for angiogenesis from arteries, however, loss of both *foxc1a* and *foxc1b* induces ectopic angiogenesis from arteries within the developing trunk through reduced Dll4/Notch signalling (Fig. 6, 7). Since *foxc1b* is not expressed in ECs within the developing zebrafish trunk, but is expressed in neighbouring somitic tissues including sclerotome (Fig. S8) (Topczewska et al., 2001b), this suggests *foxc1b* induces Dll4/Notch signalling in ECs non-cell autonomously.

In mice, non-cell autonomous anti-angiogenic functions for *Foxc1* have been described, for example, *Foxc1* expression in neural crest suppresses corneal angiogenesis via a mechanism which antagonises EC response to VEGF signalling (Seo et al., 2012). Counterintuitively, increased angiogenesis induced by *Foxc1* KO in neural crest correlated with increased corneal expression of *sVegfr1*/*sFlt1*, suggesting *Foxc1* regulates competing pro-and anti-angiogenic mechanisms (Koo and Kume, 2013; Seo et al., 2012). Similarly in zebrafish, we find context-dependent functions of *foxc1a* and *foxc1b* in suppressing angiogenesis from arteries, while promoting angiogenesis from veins. In contrast to the mouse cornea where *Foxc1* expression in neural crest cells suppresses *sFlt1* (Seo et al., 2012), *foxc1a* and *foxc1b* promote expression of anti-angiogenic *sflt1* within the DA in zebrafish (Fig. 5K, L). *sflt1* has recently been demonstrated to antagonise sprouting of veins in zebrafish (Matsuoka et al., 2016; Wild et al., 2017), and induction of *sflt1* may therefore represent an additional mechanism by which *Foxc1* limits venous angiogenesis within the developing trunk, for example, to balance its pro-angiogenic effects on SeV sprouting from the PCV (Fig. S9). Interestingly, we observed reduced expression of *kdrl* in SeAs of *foxc1a* and *foxc1a; foxc1b* double mutants, which is consistent with studies which demonstrate Foxc proteins function co-operatively with ETS transcription factors to induce endothelial gene expression and directly bind enhancers within *vegfr2/kdr* (De Val et al., 2008). However, these studies also reported reduced sprouting of intersegmental vessels (ISVs) at 24hpf following combined knockdown of *foxc1a/b* by morpholino (De Val et al., 2008), whereas our mutant analysis showed that despite reductions in *kdrl* expression following loss of *foxc1a* and *foxc1b*, SeA sprouting was not reduced, but enhanced via a Dll4/Notch dependent mechanism (Fig. 6). However, since inhibition of ISV formation was observed at very high doses of morpholino, off target effects cannot be excluded as a potential cause for these differences (De Val et al., 2008). Collectively, our data suggests that *foxc1a* and *foxc1b* control angiogenesis throughout the zebrafish vasculature by positively regulating both the pro-and anti-angiogenic inputs of the VEGF-Dll4/Notch negative feedback loop. *foxc1a* and *foxc1b* promote VEGF signalling by inducing expression of VEGF receptors within both veins and arteries. Since Notch signalling is active in arteries and actively repressed in veins (Lawson et al., 2001; You et al., 2005) *foxc1a* and *foxc1b* mediated induction of Dll4/Notch signalling serves to counteract the pro-angiogenic effects of these transcription factors in arteries but not veins. In doing so, *foxc1a/b* provide an additional level of transcriptional control to balance arterial and venous angiogenesis within developing vascular beds (Figure 7).

## Materials and Methods

### Zebrafish strains

All zebrafish were maintained according to institutional and national ethical and animal welfare guidelines. The following zebrafish lines were employed: *Tg(fli1a:EGFP)*^*y1*^ (Lawson and Weinstein, 2002), *Tg(-0.8flt1:RFP)*^*hu5333*^ (Bussmann et al., 2010), *Tg(kdrl:HRAS-mCherry-CAAX)*^*s916*^ (Hogan et al., 2009a), *Tg(flk1:EGFP-NLS)*^*zf109*^ (Zygmunt et al., 2011), *Tg(dll4in3:GFP*)^*lcr1*^ (Sacilotto et al., 2013), *Tg (hs:gal4); Tg(5xUAS-E1b:6xMYC-notch1a)* (Scheer and Campos-Ortega, 1999), *foxc1a*^*sh356*^ and *foxc1b*^*sh408*^.

### Bioinformatic analysis of *foxc1a* and *foxc1b* synteny

Orthology information for the gene of interest, plus the nearest neighbouring genes were mined from the Ensembl API (accessed October 2015). If no known orthologous gene was present in a species of interest then BLAST was used to identify the closest three genome hits with an e-value less than 1E-10. If a BLAST hit was not within a known gene model then the likelihood of an unannotated gene being present was manually analysed using EST, RNA-Seq sequences and GenScan data. However, in the majority of cases an Ensembl gene was identified and therefore orthology was determined based on synteny.

### Generation, selection and genotyping of *foxc1a* and *foxc1b* mutant alleles*foxc1a*^*sh356*^ allele

Zinc finger nucleases (ZFN) specific for *foxc1a* (ENSDARG00000091481) were generated via context dependent assembly (Sander et al., 2011) targeting the following sequence 5’-gTACCCCGCCAGCATGGCGAGGGCa-3’. For the left and right subunits, zinc fingers were added by PCR using pCS2ta3LFok1 and pCS2ta3RFok1 respectively (Ben et al., 2011) as templates, and primers listed in Supplementary Table 1. PCR products were digested with AgeI and self-ligated to generate pCS2foxc1a5-1L and pCS2foxc1a5-1R. Capped mRNA from each plasmid was generated by *in vitro* transcription and 800-1600pg mRNA injected per embryo such that embryos had an appropriate 30% rate of deformity at 24hpf. To detect potential somatic mutations, genomic DNA extracted from non-deformed embryos was amplified by PCR using foxc1a5-1F and foxc1a5-1R genotyping primers (Supplementary Table 1). Roche Titanium 454 amplicon sequencing identified 90/871 (10%) amplicon molecules included insertions or deletions at the target site. G0 adults derived from embryos injected with ZFN capped mRNA were in-crossed and G1 progeny genotyped by PCR. The *sh356* allele contains a 4bp insertion which generates a unique NsiI restriction site (Figure S3). Genotyping was performed as described previously (Wilkinson et al., 2013) followed by restriction fragment length polymorphism (RFLP) analysis. Following digestion with NsiI, the *foxc1a*^*sh356*^ allele generates fragment sizes of 119bp and 71bp in comparison to 186bp for WT *foxc1a* allele.

### *foxc1b*^*sh408*^ allele

TALENs specific for *foxc1b* (ENSDARG00000055398) were designed against the following sequence 5’-ctcgcgCATATGggccgg-3’ to target an NdeI restriction site upstream of the conserved forkhead DNA binding domain. TALENs were assembled using the Golden Gate TALEN and TAL Effector Kit (Addgene, MA, USA) (Cermak et al., 2011) to generate the pFoxc1bTal1L and pFoxc1bTal1R plasmids. Following linearisation of pFoxc1bTal1L/R with NotI, capped mRNA was generated by *in vitro* transcription and 1500pg TALEN mRNA was injected per embryo. Individual G0 embryos were tested by PCR and RFLP analysis using foxc1bTAL1F and foxc1bTAL1R (Supplementary Table 1) to identify somatic mutations which destroyed the NdeI restriction site. The progeny of TALEN injected G0 adults were incrossed and genotyped to confirm the presence of the SH408 allele, which consists of a 13bp deletion which destroys a unique NdeI restriction site (Figure S3). Following digestion with NdeI, WT *foxc1b* allele generates fragment sizes of 130bp and 98bp, whereas the *foxc1b*^*sh408*^ allele generates an undigested 215bp fragment.

### Plasmid construction and full length mRNA synthesis

*foxc1a* coding sequence from *Danio rerio foxc1a* cDNA clone IMAGE: 6789584 was cloned into pCS2+ using EcoRI and XhoI. pCS2-*sflt1* was kindly provided by Ferdinand le Noble (Krueger et al., 2011). Capped mRNA was generated using mMessage Machine SP6 Kit (Ambion).

### *In situ* probes

*foxc1a* and *foxc1b in situ* probes were generated by PCR amplification using primer sets foxc1aT3F/T7R, foxc1bT3F/T7R and IMAGE: 6789584 (foxc1a) or IMAGE: 5601888 (foxc1b) as templates. PCR products were subsequently transcribed with RNA polymerase. *xirp2a* was kindly provided by Salim Seyfried (Otten et al., 2012).

### Whole-mount *in situ* hybridisation

Whole mount colorimetric and fluorescent *in situ* hybridisation was performed as previously described (Wilkinson et al., 2012) (Thambyrajah et al., 2016). GFP expression was detected using Anti-GFP antibody (TP401; amsbio; 1:1000) alongside anti-DIG-POD antibody. Embryos were incubated with AlexaFluor-488 secondary antibody (Invitrogen; 1:500) and imaged.

### Microinjection

0.4ng *vegfc* ATG morpholino 5′-GAAAATCCAAATAAGTGCATTTTAG-3′ (Genetools) (Hogan et al., 2009a) or standard control morpholino (5’-CCTCTTACCTCAGTTACAATTTATA-3’) (Genetools) (Lee et al., 2002) were injected into one cell stage embryos.

### Heat shock induction of UAS-NICD

Heat shock was performed at 18s by incubating embryos with pre-warmed E3 at 37°C for 30mins. Embryos were maintained at 28.5°C following heatshock.

### Microscopy and image processing

Confocal images were collected using Perkin Elmer Ultraview Vox microscope and lightsheet images were performed using a ZEISS Lightsheet Z.1 microscope. Spinning disk confocal images were analysed using Volocity (PerkinElmer) and Lightsheet images were analysed with ZEN software. Images of embryos following *in situ* hybridisation was taken using a Leica M165FC and Leica DFC and imaged using Leica Application Suite software (LAS v4.3.0). Image analysis was performed using ImageJ.

### Statistical Analysis

All statistical analysis used two-tailed tests and was performed using GraphPad Prism 7. All error bars display the mean and standard deviation. P values, unless exact value is listed, are as follows: *=<0.05, **=<0.01, ***=<0.001, ****=<0.0001.

## Acknowledgements

We thank the University of Sheffield aquarium team for excellent care of zebrafish, Michael Moorhouse for technical assistance with amplicon alignments; Cecille Otten, Salim Abdelilah-Seyfried, Ferdinand le Noble and Martin Gering for probes; Yvonne Padberg, Stefan-Schulte-Merker and Wilson Clements for sharing unpublished data. This work was supported by a JG Graves Medical Research Fellowship awarded to R.N.W, Royal Society Research Grant RG120564 awarded to R.N.W, University of Sheffield Faculty Scholarship awarded to Z.J, BHF Infrastructure Grant IG/15/1/31328 awarded to T.J.A.C and R.N.W, and MRC grant MR/N020979/1 supporting T.E and M.L. The authors declare no competing interests.

---

**Supplementary Table 1.**
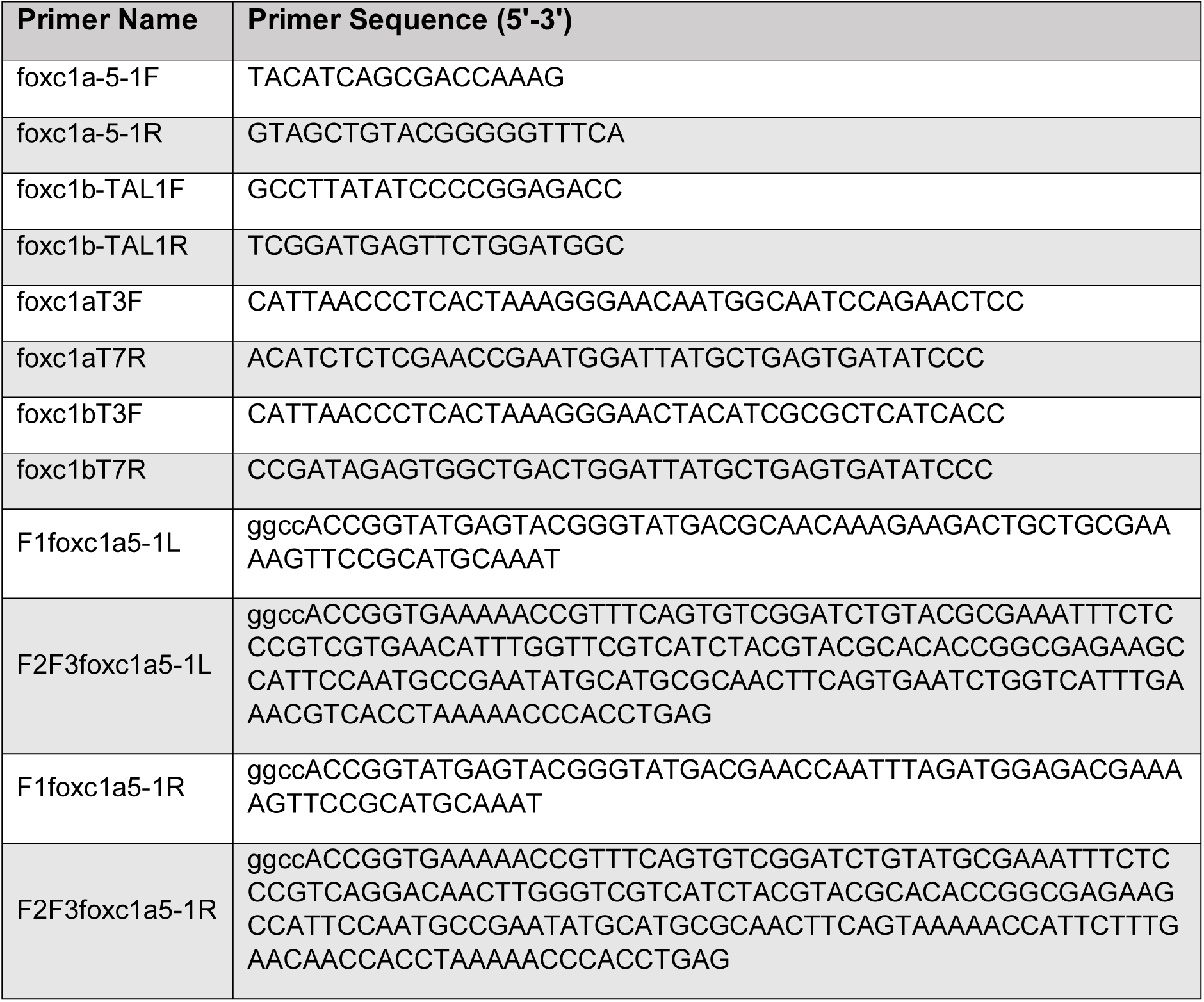

